# Proprioceptive neurons of larval *Drosophila melanogaster* show direction selective activity requiring the mechanosensory channel TMC

**DOI:** 10.1101/463216

**Authors:** Liping He, Sarun Gulyanon, Mirna Mihovilovic Skanata, Doycho Karagyozov, Ellie Heckscher, Michael Krieg, Gavriil Tsechpenakis, Marc Gershow, W. Daniel Tracey

## Abstract

*Drosophila* Transmembrane channel-like (Tmc) is a protein that functions in larval proprioception. The closely related TMC1 protein is required for mammalian hearing, and is a pore forming subunit of the hair cell mechanotransduction channel. In hair cells, TMC1 is gated by small deflections of microvilli that produce tension on extracellular tip-links that connect adjacent villi. How Tmc might be gated in larval proprioceptors, which are neurons having a morphology that is completely distinct from hair cells, is unknown. Here, we have used high-speed confocal microscopy to both measure displacements of proprioceptive sensory dendrites during larval movement, and to optically measure neural activity of the moving proprioceptors. Unexpectedly, the pattern of dendrite deformation for distinct neurons was unique and differed depending on the direction of locomotion: ddaE neuron dendrites were strongly curved by forward locomotion while the dendrites of ddaD were more strongly deformed by backward locomotion. Furthermore, GCaMP6f calcium signals recorded in the proprioceptive neurons during locomotion indicated tuning to the direction of movement. ddaE showed strong activation during forward locomotion while ddaD showed responses that were strongest during backwards locomotion. Peripheral proprioceptive neurons in animals mutant for *Tmc* showed a near complete loss of movement related calcium signals. As the strength of the responses of wild type animals was correlated with dendrite curvature, we propose that Tmc channels may be activated by membrane curvature in dendrites that are exposed to strain. Our findings begin to explain how distinct cellular systems rely on a common molecular pathway for mechanosensory responses.

## Introduction

The *Drosophila* larva is emerging as a premier model system for understanding the underlying principles of how neural circuitry is formed during development and how these circuits function to generate complex patterns of behavior. A concerted effort is underway to generate an atlas of every neuron in the larval brain and nerve cord and to assemble the complete connectome of the first instar larval central nervous system (which is composed of approximately 10,000 neurons)[1-4].

The behavioral motif that predominates larval behavior is known as peristaltic locomotion, or crawling[5]. Larvae crawl in the forward direction using rostral to caudal waves of segmental muscle contraction and they crawl backwards using an oppositely directed wave. Evidence suggests that sensory feedback is critical for the coordination of the waves of muscle contraction that occur during larval crawling. For instance, genetic silencing of specific sensory neurons dramatically impairs the procession of the waves. Two classes of arborizing multidendritic (md) sensory neurons, known as the Class I neurons (md-I) and the bipolar md neurons (md-bp) have been implicated for their critical importance[6, 7]. Silencing of either type alone causes impairments of crawling locomotion while silencing of both classes causes even more severe impairment[6]. The mild effects that come from silencing of either class alone suggests that the two classes may provide distinct parallel inputs to the brain and that the information provided by them may be partially redundant. These neurons have been proposed to send a “mission-accomplished” signal that informs the larval brain that the contraction of the muscles in a segment is complete[6]. This signal in turn is thought to facilitate the propagation of a wave neural activity into the next neuromere of the abdominal ganglion which then outputs to muscles in the next body segment as well as facilitating the termination of a contraction within a segment[6].

Although silencing the outputs of md-I neurons and md-bp neurons indicates that they are important, it remains unknown exactly how these neurons are activated during larval locomotion. In addition, how the dendrites of these cells are deformed by the stresses generated during a muscle contraction and how these associated forces are transformed into neural signals is not known. Here, we describe novel imaging preparations that have allowed us to explore these questions in semi-constrained and in freely moving animals and to investigate the molecular and mechanical mechanisms involved with the proprioceptive responses.

Surprisingly, we find that the Class I neurons show preferred responses to movement direction. In addition, we find that the putative mechanosensory channel TMC is required for peripheral responses during both forward and backward movement.

## Results

There are three md-I neurons in each body segment with dendrites that run along the basolateral surface of epidermal cells: a single ventral neuron (vp-da) and two dorsal neurons (ddaE and ddaD)[8]. The “receptive fields” innervated by the dendrites of these neurons are non-overlapping[8]. The ddaE neurites are localized to the posterior compartment of each body segment. The primary dendrite of ddaE projects dorsally and its secondary and tertiary dendrites are directed towards the posterior of each body segment (Figure 1A). In contrast, the ddaD neuron innervates the anterior compartment with an apparent mirror-image symmetry to ddaE (Figure 1A). As with ddaE, ddaD has a dorsal directed primary dendrite but it’s secondary and tertiary dendrites project anteriorly (Figure 1A). The vp-da neuron innervates a posterior compartment on the ventral side (not shown). It has a single dorsally directed primary branch and secondary dendrites that are directed both anteriorly and posteriorly[8]. In this study, we investigate ddaE and ddaD.

**Figure 1:**
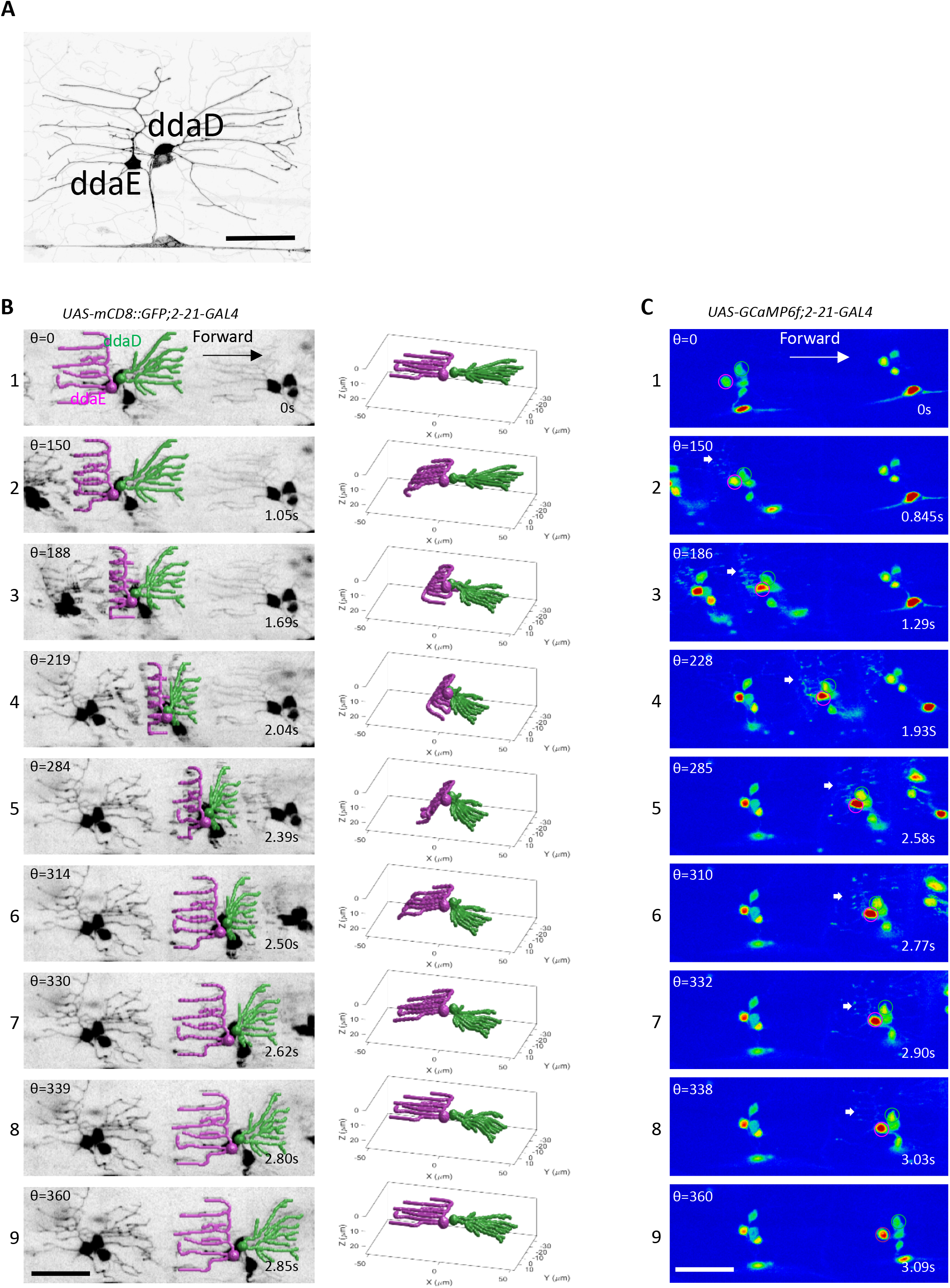
Dendrite deformation and GCaMP6f activation patterns in larval forward locomotion. **A)** Dendrite morphology of class I da neurons ddaE and ddaD visualized by expression of mCD8::GFP driven by 2-21-GAL4 in first instar larvae. **B)** Dendrite deformation pattern of ddaE and ddaD during forward locomotion. Left panel shows individual time points of a maximum intensity projection from a volumetric time series. Right panel depicts the model of dendritic architecture reconstituted by a computer vision framework for neurite tracing from the volume shown in the left panel. ddaE and ddaD are colored in magenta and green respectively. **C)** GCaMP6f activation pattern of ddaE and ddaD during forward locomotion. The images at the nearly similar phase of segmental contraction cycle shown in the left panel of B) are selected to show the activation of GCaMP6f. ddaE and ddaD are marked with magenta and green circles respectively. Arrows point to activated GCaMP6f in ddaE dendrites. Maximimum intensity projections of confocal Z time series are shown in A and C. All images are show as dorsal side up and anterior on the right. Scale bar is 40μm.

Our first goal in providing a better understanding of how these neurons function in proprioception was to observe and describe the types of movement related deformations that can be seen in the sensory dendrites during larval locomotion. To do so, we placed larvae in three-sided straight agarose channels that were filled with water and placed upon a cover glass [5]. This arrangement restricted the larval motion to a straight line through the agarose channel and this allowed us to observe its movement through the cover glass on an inverted confocal microscope.

A high-speed microscope used in our study was equipped with a piezo driven objective lens. The former allowed for rapid acquisition of XY optical sections (at over 200 Hz) and the latter allowed for rapid changes of the focal plane and high-speed acquisition of three-dimensional Z-stacks (9-15 volumes per second). This configuration was essential for our study because the dendrites and cell soma could be captured at high resolution even though the objects of interest were moving in the XY and Z axes during the acquisition of our recordings. The rapid scan rate allowed us to observe the objects of interest in focus because we could capture them somewhere within the deepest and shallowest section of our Z-stacks throughout the periods of observation.

### Forward and backward locomotion uniquely deform Class I dendrites

We first observed the neurons in larvae that were moving across the field of view by labeling their cell bodies and dendrites with the fluorescent membrane marker mCD8::GFP. Prior to the initiation of a segmental muscle contraction, the dendrites of the md-I neurons ddaD and ddaE were seen as they are normally depicted when stationary in the many studies that have used these dendrites for the study of dendrite morphogenesis. In the relaxed body segment the dendrites lay relatively flat beneath the epidermis (Figure 1A,B1, Figure 2A). But during locomotion the dendrites can be seen to bend as they are exposed to the stresses exerted by muscles contracting within the segment (Figure 1B1-B9, Videos S1,S2). During forward locomotion, as the wave of segmental muscle contraction proceeded from posterior to anterior, the dendrites of ddaE were the first to be deformed by the contraction forces within a segment (Figure 1B, Videos S1,S2). The dendrites could be seen to bend toward the interior of the larva in a pattern that began at the distal tips (Figure 1B, Videos S1,S2). Then, as the contraction proceeded, the deformation of the dendrites travelled from the distal dendrites towards the proximal dendrites and the more proximal dendrites were gradually pulled deeper within the deformed body segment (Figure 1B2,B3, Videos S1,S2). This pattern of forces caused the distal ends that were nearest to the segmental boundary to lay deepest within the imaging plane (Figure 1B3,B4, Videos S1,S2). As the segmental contraction initiated, the dendrites of ddaE showed clear deformation, even though the dendrites of ddaD remained initially stable and relatively unmoved by the muscle driven stresses. Thus, the ddaE neuron dendrites showed an earlier deformation during the muscle contraction cycle relative to ddaD. Surprisingly, the initial stability of the ddaD dendrites suggested an asymmetry in the stresses applied to dendrites in the posterior and anterior compartments during a segmental contraction. Thus, unlike the depiction of previous studies, a contracting segment does not simply compress the epidermis like an accordion. Rather, our imaging of the strain field suggests that the strain travels as a wave from one side of the segment to the other.

**Figure 2:**
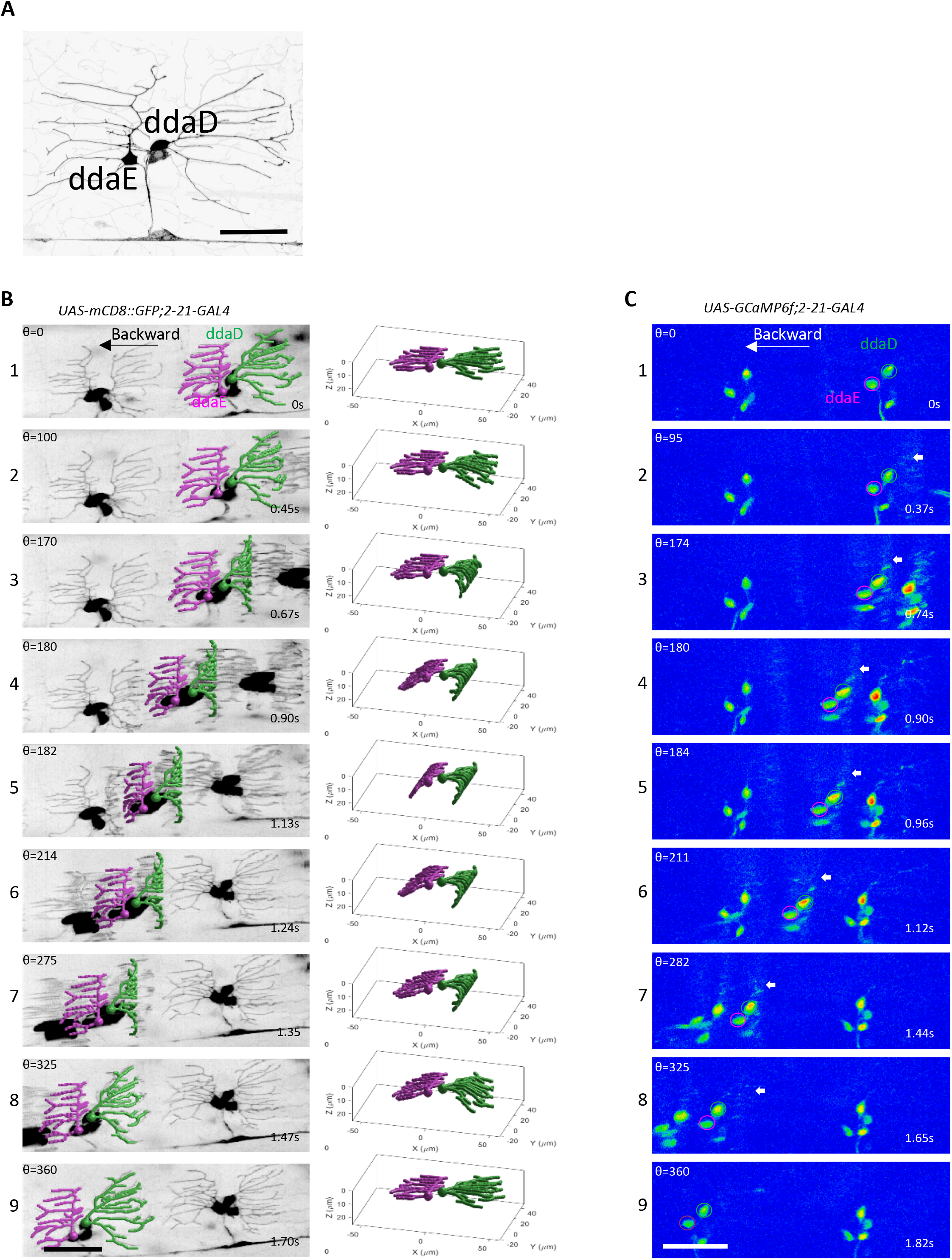
Dendrite deformation and GCaMP6f activation patterns in larval backward locomotion. **A)** Dendrite morphology of class I da neurons ddaE and ddaD visualized by expression of mCD8::GFP driven by 2-21-GAL4 in first instar larvae. **B)** Dendrite deformation pattern of ddaE and ddaD during backward locomotion. Left panel shows individual time points of a maximum intensity projection from a volumetric time series. Right panel depicts the model of dendritic architecture reconstituted by a computer vision framework for neurite tracing from the volume shown in the left panel. **C)** GCaMP6f activation patterns of ddaE and ddaD dendrites in backward locomotion. Representative images at the nearly similar phase of segmental contraction cycle shown in the left panel of B) are selected to show the activation of GCaMP6f. ddaE and ddaD are marked with magenta and green circles respectively. Arrows point to activated GCaMP6f in ddaD dendrites. Maximum intensity projections of confocal Z time series are shown in A and C.All images are show as dorsal side up and anterior on the right. Scale bar is 40μm.

Eventually, as the muscle contraction strengthened, forward momentum of the neuronal cell bodies could be seen. It was around this point that the dendrites of ddaD began to be deformed by the forces of the segmental contraction (Figure 1B3, Videos S1,S2). In ddaD, the earliest deformations were seen in proximal dendrites while the distal tips remained relatively stationary. As the muscle contraction progressed, the wave of dendrite deformation traversed from proximal to distal in the dendritic tree of ddaD (Figure 1B3-B7, Video S1,S2). Thus, the dendrite displacement for ddaD and ddaE differed in at least two ways during a segmental contraction. First, dendrites of the two neurons were deformed at different phases of the contraction cycle (with ddaE showing signs of deformation nearer to the beginning of the cycle and ddaD displacements occurring later). Second, the dendrites of ddaE showed deformations that began distally and spread proximally while the dendrites of ddaD deformed in a proximal to distal progression.

During backwards locomotion, the pattern of dendrite deformation that we saw was not simply the forward locomotion pattern played in reverse (Figure 2B1-9, Videos S3,S4). It was the mirror image. As a segment began its muscle contraction, it was the dendrites of ddaD that showed the early inward bending pattern beginning at the distal tips of dendrites while the dendrites of ddaE remained relatively stationary (Figure 2, Videos S3,S4). Then, as the muscles contracted further the ddaE dendrites showed a later pattern of deformation (Figures 2B2-B3, Videos S3,S4). During backwards locomotion it was the dendrites of ddaD that showed a distal to proximal progression of deformation while the ddaE dendrites showed the proximal to distal deformation pattern (Figures 2B4-B6, Videos S3,S4).

### Class I neurons show preferred response to movement direction

To determine how these patterns of dendritic deformation were converted into neuronal signals, we next observed larvae that were expressing the genetically encoded Calcium sensor GCaMP6f in the moving larval preparation. In stationary larvae, the baseline fluorescence for GCaMP6f in the dendrites was low but fluorescence in the cell bodies could be detected. During locomotion the dendrites of the Class I neurons showed striking increases in fluorescence (Figure 1C1-9, Video S5). This dendritic signal evolved into a strong increase of somatic fluorescence as the muscle contraction proceeded. The presence of detectable baseline fluorescence in the cell soma enabled tracking of the cells and quantification throughout the experiment using a custom build Matlab program.

As with the patterns of dendrite deformation that we observed, the calcium responses of ddaE and ddaD were distinct during locomotion. The peak fluorescence increase for ddaE (measured at the soma) during forward locomotion was substantially higher than the increase that was seen for ddaD (Figure 1C, Figure 3A, Video S5). Thus, the ddaE neuron is strongly activated by the forces of forward larval locomotion while the ddaD neuron is less strongly activated. In contrast, during backwards locomotion the peak fluorescence increase for ddaD was substantially higher than that of ddaE (Figure 2C1-9, Figure 3A, Video S6). The ddaD neuron therefore is more strongly tuned to respond to contractions in a segment during backwards locomotion.

**Figure 3:**
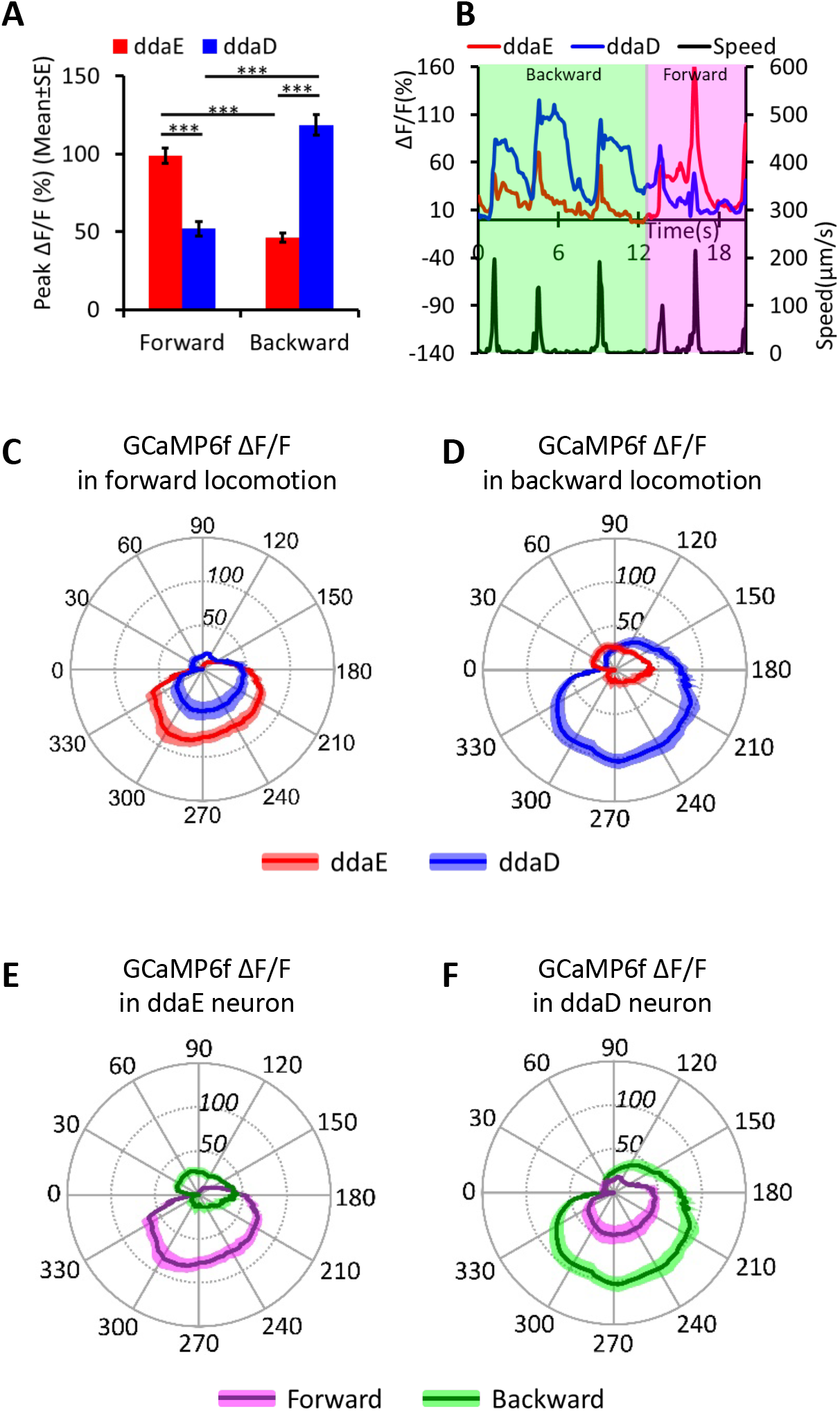
Preferential activation of class I da neurons (ddaE and ddaD) during larval forward and backward locomotion. **A)** Comparison of peak ΔF/F in ddaE and ddaD during forward and backward locomotion. *** P<0.001; Student’s *t*-test. (n=25-39 forward movements), (n=25 backward movements). **B)** Trace of GCaMP6f ΔF/F for ddaE and ddaD neuron of the same segment during three consecutive waves of backward and three waves of forward locomotion. The bottom frame shows the speed of the ddaE neuron movement throughout the time series. **C-D)** Comparison of mean ΔF/F for ddaE and ddaD in the segmental contraction cycle during forward (n=12) and backward (n=9) locomotion. **E-F)** Comparison of mean ΔF/F in ddaE (E) and ddaD (F) in forward and backward locomotion in a segmental contraction cycle. **C-F)** Position along the radial axis represents ΔF/F (percent change). The phase angle of the segmental contraction cycle depicts the distance between neurons of adjacent segments, which is decreasing from 0-180 degrees and increasing from 180-360 degrees. Darker line indicates mean ΔF/F and the lighter colored shading denotes standard error of mean.

These direction specific responses are vividly illustrated in recordings where ddaE and ddaD within a single segment could be observed continuously when an animal reversed direction during the recording. For instance, as depicted in Figure 3B, an animal was recorded as it moved backwards for three cycles of contraction and then switched to forwards locomotion for another three cycles. Strong increases in calcium signals are seen in ddaD during the first three cycles of backwards locomotion and relatively weak signals are seen in the same neuron during the subsequent three cycles of forward locomotion (Figure 3B, Video S7). In contrast, the ddaE neuron in this same segment shows relatively weak increases in calcium responses during the initial period of backwards locomotion but then shows a strong response during the waves of forward locomotion (Figure 3B, Video S7). The fact that the cell bodies of ddaE and ddaD neurons are side-by-side in this and all of our recordings, but yet they show very distinct responses to movement, is strong evidence that the calcium signals observed are genuine and not a result of a motion artifact of the recordings. In addition, the Ca^++^ transients revealed by GCaMP6f increases often outlasted the period of movement by the cell body. Consistent with the proposal that the signals recorded relate to neuronal activity rather than movement artifact, control recordings made from animals expressing mCD8::GFP in the class neurons showed relatively little change in fluorescence during movement (Figure S1, Video S8). In addition, as described below, the directional selective properties of the ddaE and ddaD neurons were similar in a freely moving larval preparation that allowed for ratiometric measurements (a rigorous correction for movement) with a calcium insensitive red fluorescent protein.

### Proprioceptive responses relate to the phase of the segmental contraction cycle

Although the changes in GCaMP6f fluorescence that occurred during our calcium imaging could be extracted from the recordings, understanding how the responses related to the phases of the muscle contraction cycle required a method for comparing across animals and neurons. We could not simply compare time courses of the responses because each of the animals moved at different speeds during our experiments and the animals also slightly varied in size. We overcame these difficulties by plotting the calcium responses of the Class I neurons as a function of the contraction cycle that could be measured between two adjacent segments. We used the position of the cell bodies of neurons in adjacent segments as fiduciary landmarks, as this gave us a readout of the phase of the segmental contraction cycle. As a body segment contracts, the distance (D) from the neuronal cell bodies contained within the segment to the cell bodies in an adjacent segment is shortened, and when the contraction relaxes, D increases back to the initial maximum (Figure S2). Therefore, we could use the distance between class I neuron cell soma in neighboring segments as a readout of the stride. The D we measured between cell bodies in adjacent segments was converted to polar angle coordinates (theta), and the change in GCaMP6f fluorescence was plotted on the radial axes of these coordinates (note that theta is also indicated for representative images in Figures 1B,C and Figure 2 B,C). In these plots, the period of shortening D between the cell soma traverses clockwise from 0 to 180 degrees and the subsequent period of increasing D continues from 180 to 360 degrees (see Methods, Figure S2). When the data were plotted in this way it was clear that reproducible and consistent patterns of calcium activation were present across animals and that the pattern of measured activity related to the cycle of the segmental contraction.

During forward locomotion a steep rise in ddaE activity is seen as its cell body is approached by ddaE in the next more posterior segment (Figure 3C,E). This activity peaks when D is near a minimum (at a theta of approximately 180-210 degrees) and is maintained as the distance between the cell soma in adjacent segments relaxes. The activity does not rapidly decay until (approximately 300-330 degrees) shortly before relaxation to the maximum distance between the cells. As with the peak responses, the calcium responses of ddaD, were significantly weaker throughout the entire contraction cycle of forward locomotion (Figure 3C,F). Interestingly, during backward locomotion the pattern of activity reflected in the segmental contraction cycle appears identical to what is seen during forward locomotion except that the identity of the neurons is switched. In this case it is ddaD that shows the increasing signals as it is approached by the cell body of the next more anterior segment and the ddaE responses were significantly smaller (Figure 3 D).

### Calcium responses of Class I neurons in a freely moving larval preparation

The movement of larvae through the agarose channels on the high-speed confocal microscope enabled us to observe dendritic features and to measure calcium responses. But this preparation limited our analysis to larvae constrained to crawl in a straight line. Furthermore, because our confocal system was not enabled to continuously track moving neurons and could only record from a single fluorophore, we were able to record only for short epochs and could not use ratiometric techniques to completely correct for motion-related artifacts that were unrelated to neural activity. Therefore, to confirm our finding that ddaE and ddaD neurons are selectively active during forward and backward crawling and to seek additional insight into the dynamics, we recorded from these neurons using a recently described tracking two-photon fluorescence microscope[9]. This microscope follows the cell bodies of targeted neurons using real-time hardware feedback while simultaneously monitoring both GCaMP6f, the calcium indicator, and hexameric mCherry, a stable reference indicator. To track neurons, the microscope scans the focal spot rapidly within the soma, allowing us to continuously monitor the neuronal activity via the ratio of GCaMP6f to mCherry fluorescence, but not to form images.

The microscope uses sub-millisecond feedback to the scanning mirrors to follow neurons within the objective field of view and slower feedback to the stage which keep the neurons centered under the objective. Finally, a low magnification infrared camera records the larva’s behavior. Using this microscope, we made simultaneous recordings for up to 20 minutes from ddaE and ddaD in a second instar larva exploring an agar coated microfluidic arena (Figure 4A-B, Video S9). We observed spontaneous transitions between forward and backward crawling and found that as with our confocal system, ddaE and ddaD were robustly active during forward and backward crawling respectively, while remaining relatively silent during the opposite phase of motion (Figure 4A-B, Video S9). Both neurons encoded movement with activity whose peak approximately coincided with the peak velocity of the cell body, and both showed sustained elevations of calcium following the offset of motion. When we examined the fluorescence signal from the stable red indicator, we found very little variation with motion, and we did not observe the neuron-specific directional encoding we found in the activity.

**Figure 4:**
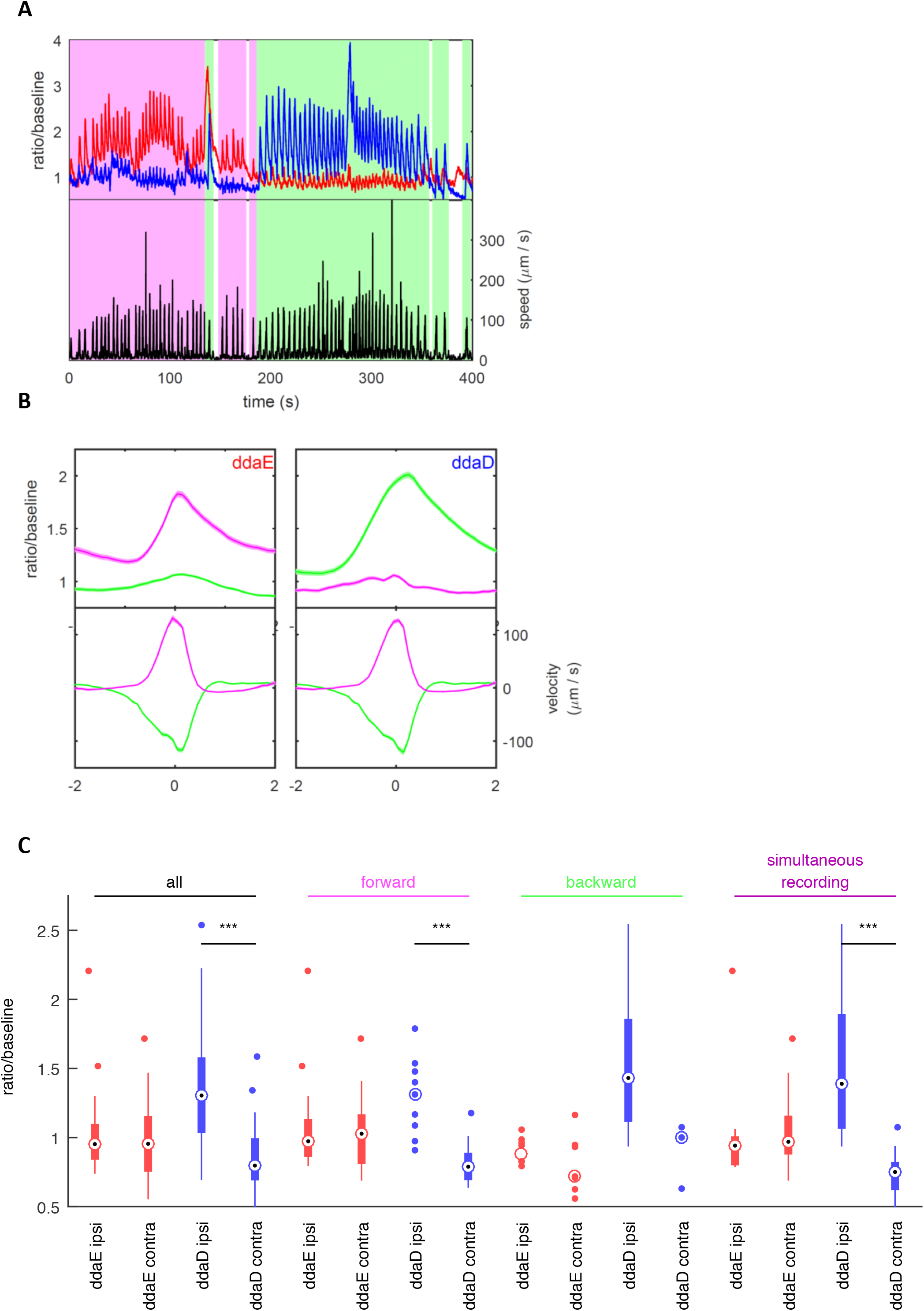
Relationship between neuron activity and locomotion in freely moving larvae. Representative traces from simultaneous recording of ddaE and ddaD during larval locomotion obtained using a two-photon tracking microscope. **A)** Activity (top panel) and instantaneous speed (bottom panel) during a 400 s period completed with backward and forward locomotion bouts. Shading indicates behavioral state (magenta = forward crawling, green = backward crawling). Activity (red = ddaE, blue = ddaD) is measured as a ratio of GCaMP6f fluorescence to mCherry fluorescence divided by a baseline ratio fit as a simple exponential to the entire data set to account for differential rates of photobleaching. Shaded region surrounding activity traces represents uncertainty in the measurement determined as an output parameter of MLE fit. **B)** Activity and velocity aligned to time within a bout. A bout is defined to be a period of rapid movement. Velocity is defined as the movement of the neuron relative to the position of the tail. Forward bouts (magenta) have positive velocity (away from the tail) and backward bouts (green) have negative velocity. The time axis is aligned so that the fastest movement is at t = 0. No measurements of activity were used to detect bouts or align the time axis. Shaded regions represent mean +/- standard error of the mean. Left panels show ddaE and right panels show ddaD (n = 182 forward bouts, both neurons; n = 224 backward bouts, both neurons). **C)** Activation of dorsal neurons during body bends. Box and whisker plot showing median, 25^th^ and 75^th^ percentiles, range of data, and outliers for activity at moment of maximum body bend, as determined by visual inspection of raw behavioral video. When a group is comprised of less than 10 data points, all points are shown. Ipsi indicates a bend toward the neuron (body wall is convex at location of neuron); contra indicates a bend away from the neuron (body wall is concave at location of neuron). Groupings: *all* – all sampled body bends of the given type; *forward* – all sampled body bends in which the larva was crawling forward before and after the bend; *backward* – all sampled body bends in which the larva was crawling forward before and after the bend; *simultaneous recording –* subset of bends measured when tracking both ddaD and ddaE simultaneously. *** at p < 0.001, rejects the hypothesis that both groups are drawn from the same random normal distribution, using two-sample t-test.

Despite the difference in microscope technology, larval stage, and microfluidic device, the results from the tracking and confocal microscopes agree quantitatively and qualitatively, making it extremely likely that these results reflect genuine neuronal activity. Additionally, because the tracking microscope records for sustained periods of motion, using this preparation we observe that not only do the magnitudes of the activity peaks encode direction, but that the baseline calcium levels are also higher in ddaE during forward movement and in ddaD during backwards crawling (Figure 4A,B).

Unlike the confocal preparation, which constrained larvae to crawl in a straight line, the tracking microscope followed larvae through an open arena in which they were able to bend their bodies and change direction. We wondered whether and how activity in the dorsal neurons might encode these postural changes. Automatic registration of larval posture using machine vision proved difficult in the videos taken under the tracking microscope. We therefore hand-selected approximately 100 bending events from recordings of larval behavior. We made these selections blind to the activity of the neuron(s) being tracked and included recordings in which only ddaD or only ddaE was tracked, as well as recordings in which two neurons were tracked simultaneously.

We found that ddaD activity increased when the larva’s body bent ipsilaterally (ie. bent towards the side of the body containing the neuron) compared to when the larva bent away from the side containing the cell body (ie. contralateral bend) (Figure 4C, Figure S4). Surprisingly, we observed no difference in ddaE activity between ipsilateral and contralateral bends (Figure 4C). We found this same pattern whether the larva bent while crawling forward or backward although during backward crawling the difference in ddaD activity was not statistically significant due to a low number of contralateral exemplars(Figure 4C). Importantly, when we considered only the experiments in which both ddaD and ddaE were tracked simultaneously, we again found ddaD alone encoded the bend direction. Therefore, it appears that ddaD uniquely responds to bends in which the animal has turned towards the side of the body that contains it. When the larva turns towards the left, the left ddaD neurons respond. When the larva turns to the right, ddaD neurons on the right side of the body respond.

### Dendrite Curvature and Proprioceptor Responses

How does the dendrite deformation that we recorded in experiments in neurons expressing mCD8::GFP relate to the physiological responses? To investigate this relationship, we plotted dendrite curvature onto polar plots again using the distance between cell soma in adjacent segments as a readout of the contraction cycle. Dendrite curvature was estimated using a machine vision “accordion model” (see Methods) and plotted in radians. In figure 5A, it can be seen that during forward locomotion the dendrites of ddaE showed an increase in curvature at an earlier phase in the contraction cycle relative to ddaD. In addition, during forward locomotion the magnitude of ddaE dendrite curvature exceeds that of ddaD. In Figure 5B, the pattern of curvature for reverse locomotion is shown and it can be seen that in this case ddaD dendrites curved earlier and with a greater magnitude relative to ddaE.

**Figure 5:**
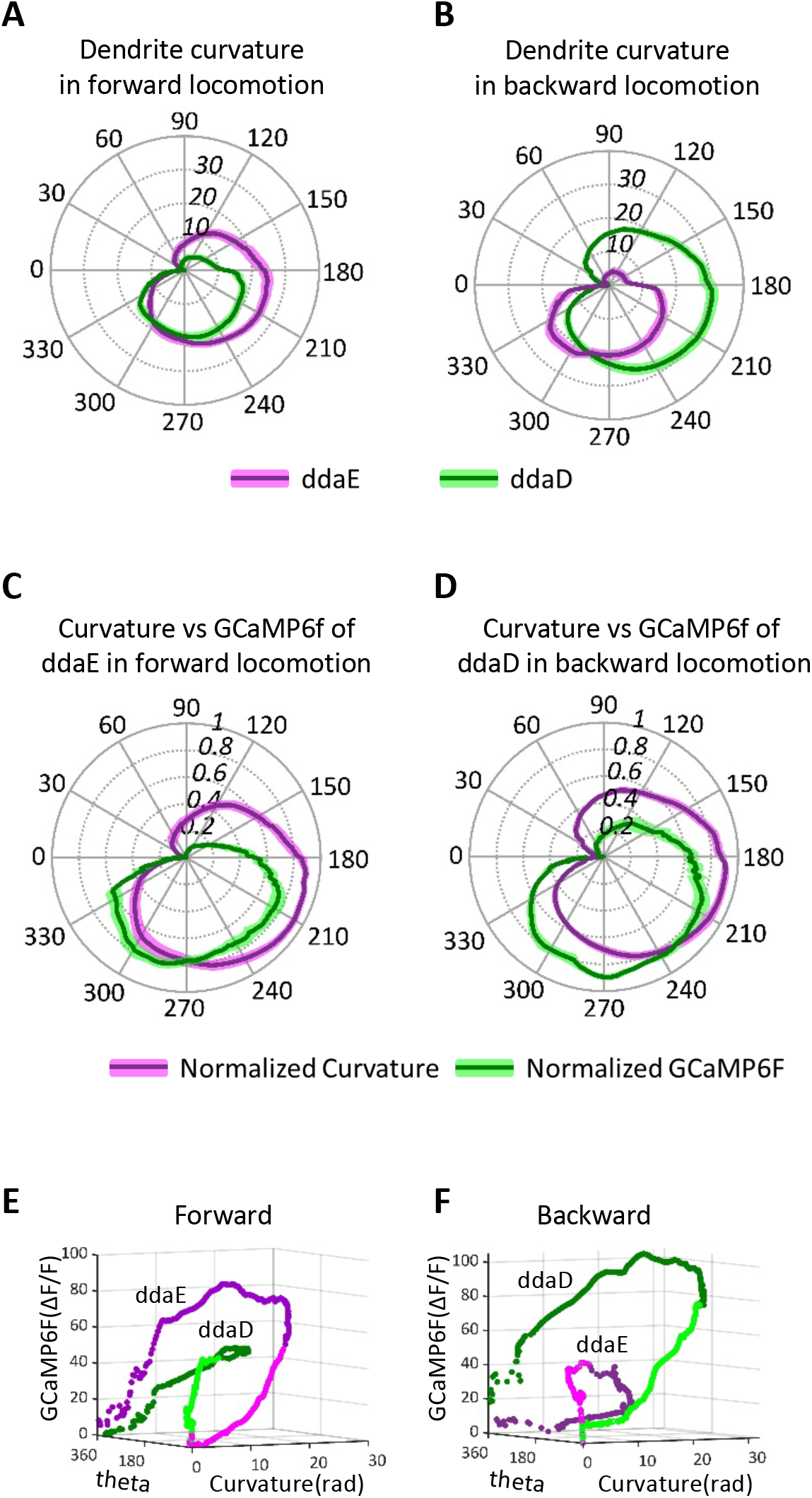
Comparison of dendrite curvature and GCaMP6f activation during larval locomotion. **A,B)** Absolute dendrite curvature of ddaE and ddaD neurons plotted versus phase of the segmental contraction cycle. Position along the radial axis represents absolute curvature. The phase angle of the segmental contraction cycle depicts the distance between adjacent neurons, which is decreasing from 0-180 degrees and increasing from 180-360 degrees. Black line represents mean of curvature (n=11 for forward locomotion; n=8 for backward locomotion), colored shading represents standard error of mean. Overall the curvature of ddaE is larger than that of ddaD during forward locomotion and vice versa during backward locomotion. **C,D)** Normalized dendrite curvature and GCaMP6f signals are plotted on the same polar coordinates shows that dendrite curvature precedes the rise in GCaMP signal and decreasing of curvature anticipates the decline of GCaMP6f signals. **E,F)** Change in GCaMP6f (z axis), curvature of dendrites (x-axis), and theta (y-axis) for ddaE and ddaD during forward and backward locomotion respectively. The ddaE neuron is shown in magenta and ddaD is shown in green. Lighter magenta and green represent phase from 0-180 degrees while darker magenta and green represent phase from 181-360 degrees.

Interestingly, when normalized dendrite curvature and change in GCaMP fluorescence are plotted on the same polar coordinates it could be seen that the curvature of dendrites occurs at an earlier phase of the segmental contraction cycle relative to the rise in GCaMP6f fluorescence measured at the soma (Figure 5C,D). Although dendrite curvature and GCaMP measurements were not made in the same animals, the data suggest that dendrite bending in Class I neurons occurs upstream of the physiological responses to movement. This further suggests that the transduction machinery for sensing locomotion may be localized to the dendrites of these cells. The same data plotted without normalization further suggest a relationship between the degree of dendrite curvature that occurs in the segment contraction cycle and the strength of the neuronal response such that higher curvature is correlated with stronger GCaMP6f signals (Figure 5E,F). Thus, we propose a mechanism where dendrite curvature may drive the activation of these proprioceptors. The curvature of these dendrites might generate tension or compression in the dendritic membrane and/or cytoskeleton[10] which in turn activates a mechanosensitive channel that resides in the dendrites of these cells.

#### Molecular mechanisms of peripheral proprioceptive responses

A recent study has proposed a role for a protein encoded by the *Transmembrane channel-like* (*Tmc*)) gene as a candidate mechanoreceptor molecule that might mediate the proprioceptive responses of the class I da neurons[11]. Several observations are consistent with this hypothesis. First, the TMC protein is localized to Class I dendrites and larvae that are mutant for *Tmc* show defective and uncoordinated locomotion behavior[11]. In addition, transcriptional reporters for *Tmc* have been found to be expressed in the class I da neurons[11, 12]. Finally, calcium imaging preparations have shown that the <u>central projections</u> of class I da neurons show reduced responses to artificially applied mechanical manipulation of the larval body[11]. However, at the level of the physiological mechanism, it is unknown if *Tmc* is required in the periphery for sensory responses to idiothetic movements, or if it is solely required for electrical propagation of signals from the periphery that result from externally applied forces.

Thus, we investigated the class I GCaMP6f responses of animals that were mutant for *Tmc* in our confocal preparation. Remarkably, in homozygous *Tmc^1^* mutant animals, movement related increases in GCaMP6f were nearly eliminated (Figure 6A-B, Figure 7A-F, Videos S10 and S11). lnterestingly, while the GCaMP6f signals in the class I dendritic arbors was often seen grow brighter during wild type locomotion (highlighted by arrows in Figures 1 and 2) the dendrites of *Tmc^1^* mutant animals showed no observable increases and remained quiescent (Figure 6). The lack of an observable movement related signal in the soma (and in the sensory dendrites) suggests that the TMC protein is critical for direction selective mechanosensory responses. This further highlights the likely role of this molecule as a candidate sensor for the forces imposed on the proprioceptive dendrites during larval locomotion.

**Figure 6:**
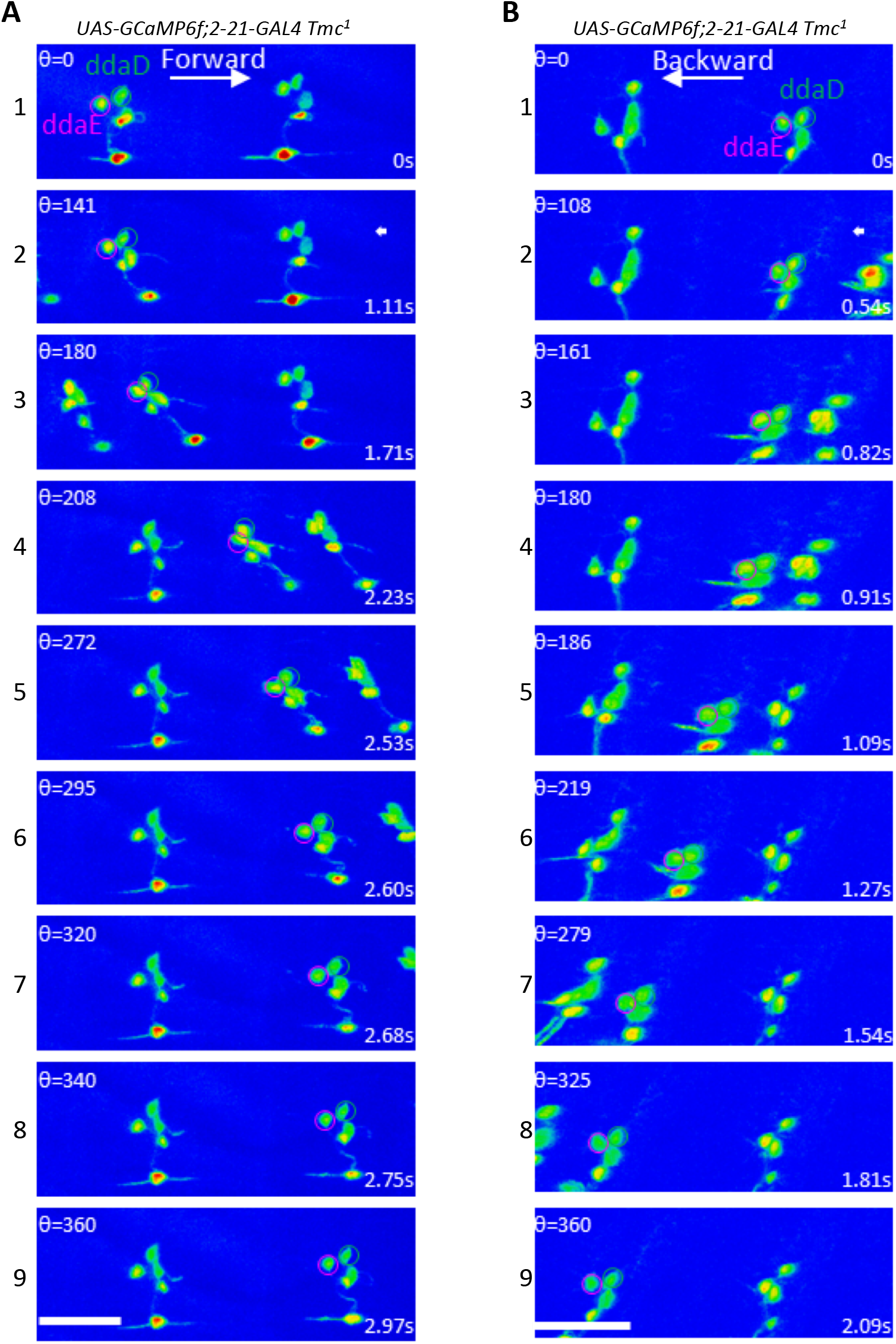
Representative still images from *GCaMP6f* recordings on *Tmc^1^* mutant larvae (*w;UAS-GCaMP6f; 2-21-GAL4 Tmc*^1^). Maximum intensity projections of confocal Z time series are shown in A and B. In all panels the ddaE cell soma is encircled in magenta and ddaD is encircled in green. Each panel indicates the value for theta and the time stamp for the relevant image. **A.)** Still images of a larva undergoing forward locomotion, note that there is little change in fluorescence intensity of ddaE. **B.)** Still images of a larva undergoing backward locomotion, note that there is little change in fluorescence intensity in ddaD.

**Figure 7:**
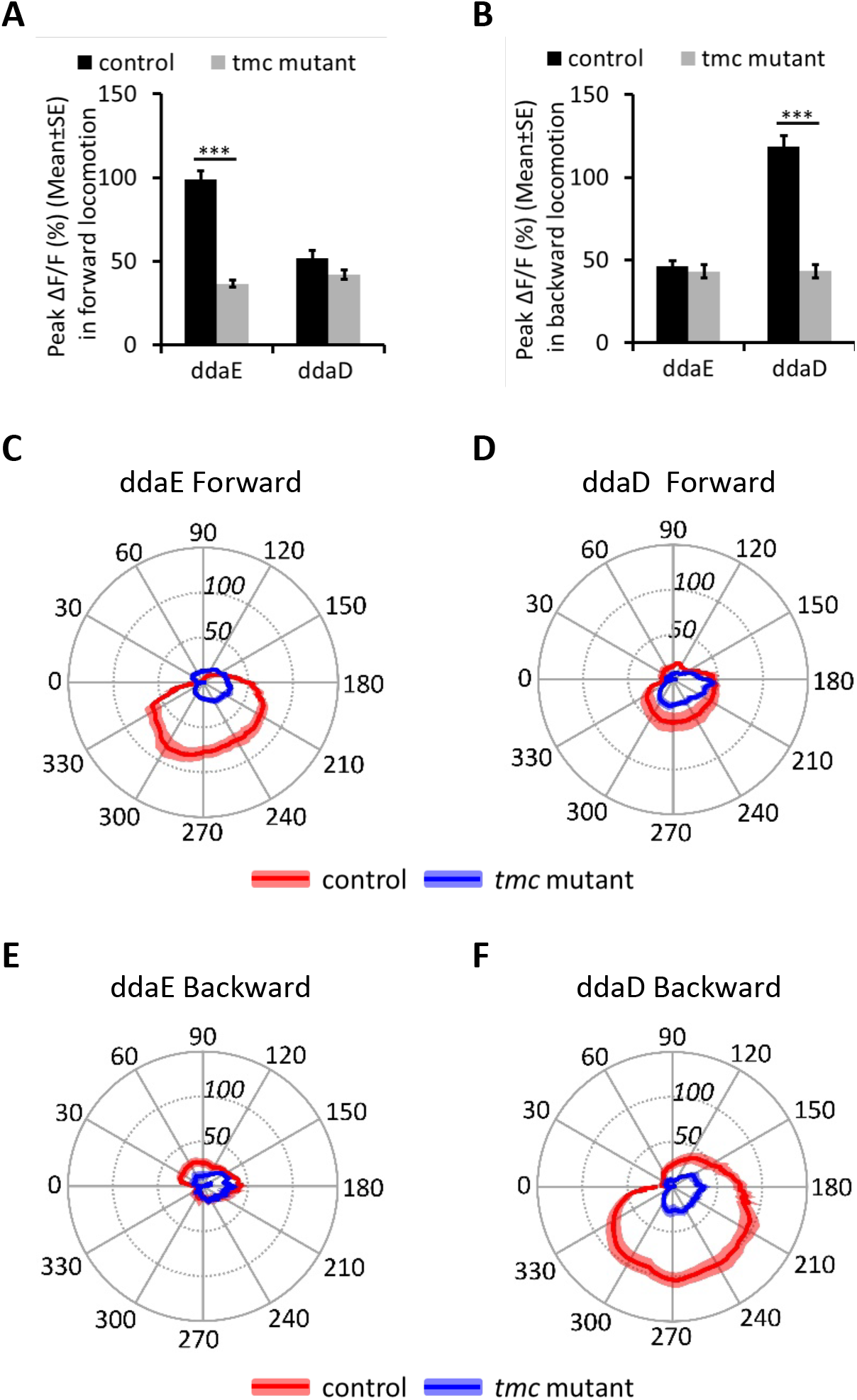
TMC is required for idiothetic activation of ddaE and ddaD in larval forward and backward locomotion. **A)** Comparison of peak ΔF/F (%) in ddaE and ddaD neurons in control (n=39,35) and *Tmc* mutant (n=39,37) in larval forward locomotion. **B)** Comparison of peak ΔF/F (%) in ddaE and ddaD in control (n=25) and *Tmc* mutant (n=15) in larval backward locomotion. **C)** Comparison of mean ΔF/F (%) in ddaE of control (n=12) and *Tmc* mutant (n=11) in larval forward locomotion in a muscle contraction cycle. **D)** Comparison of mean ΔF/F (%) in ddaD of control (n=12) and *Tmc* mutant (n=11) in larval forward locomotion in a muscle contraction cycle. **E)** Comparison of mean ΔF/F (%) in ddaE of control (n=7) and *Tmc* mutant (n=6) in larval backward locomotion in a muscle contraction cycle. **F)** Comparison of mean ΔF/F (%) in ddaD of control (n=9) and *Tmc* mutant (n=7) in larval backward locomotion in a muscle contraction cycle. Control: *UAS-GCaMP6f;2-21-GAL4*; *Tmc* mutant: *UAS-GCaMP6f;2-21-GAL4 Tmc^1^*. ***P<0.001; Students t-test. Position of the radial axis in panels C-F represents ΔF/F(%). Black line represents the mean values and colored shading denotes standard error of mean. n is the number of neurons examined.

## Discussion

For stimuli in motion, sensory systems must encode the direction of movement. This is perhaps best studied in the visual system, where neurons in the vertebrate and invertebrate retina are activated by moving edges in a visual scene[13]. In the retina, specific neurons are tuned to be activated by stimuli moving in a preferred direction but are inhibited by stimuli with non-preferred motion. More poorly understood is how mechanosensory systems might encode the direction of movement. Nevertheless, “direction selectivity” has been observed in several mechanosensory systems. In the best understood example, the hair cells of the inner ear show a preferred mechanosensory response when the actin rich bundles of stereocilia are displaced towards the microvilli on the taller side of the bundle[14]. Another example is found in the neurons that innervate the mechanosensory bristles of adult *Drosophila.* These neurons are activated by forces that displace the bristle towards the body but not by displacements away from the body[15]. Low threshold mechanoreceptors with lanceolate endings that innervate hair follicles in the mouse respond preferentially to deflection of hairs in the caudal to rostral direction[16].

Here, we have discovered another example of preferred directional mechanosensory responses in identified non-ciliated sensory neurons of the *Drosophila* larva. The ddaE neuron shows preferential responses to forward locomotion while the ddaD neuron responds preferentially to backward locomotion. Interestingly, the molecular basis of these mechanosensory responses depends on the *Drosophila Tmc* gene which encodes a putative ion channel gene that is homologous to a pore forming subunit of the mechanotransduction channel of mammalian hearing (TMC1)[17, 18].

In the hair cell, direction selectivity is an emergent property of the actin rich bundle of stereovilli. The villi possess extracellular tip-links which transmit tugging forces to the mechanosensory channels that are localized near the tips of the actin bundles[19]. The tip-link tension that is needed for mechanosensory channel gating is generated when the bundle is deflected towards the tallest side but not when deflected towards the shortest side. A dimeric TMC1 protein complex comprises an ion channel that may be activated by the tugging forces of the tip-link[17, 18]. It is remarkable that *Drosophila* proprioceptive neurons, which bear no apparent structural resemblance to the inner ear hair cell, rely on a homologous gene (*Tmc*) for mechanosensory responses that are direction sensitive. These observations raise interesting questions for future study. How can *Tmc* family channel members function for mechanosensation in such structurally distinct cells as Class I neurons and hair cells? Do Class I neurons possess extracellular or intracellular links that are involved in activating the TMC channels? If not, it may be that membrane curvature or tension alone is an important feature for the activation TMC channels. The latter idea is consistent with proposed models for activation of mechanosensory transduction channels via the forces imposed on them by the plasma membrane[20, 21]. An additional question that comes from our studies underlies finding the mechanism that generates the preferred direction responses of the Class I neurons. We envision several potential possibilities that are not mutually exclusive. The first possibility is that the direction preference is entirely explained by the magnitude of dendrite curvature that occurs in the different neurons during forward and backward movement. Our estimates of dendrite curvature were found to be higher in ddaE relative to ddaD during forward locomotion and higher in ddaD than in ddaE during backwards locomotion. Thus, in our experiments the degree of curvature was correlated with the strength of the calcium signals that we observed in the different neurons during movement. Although the total curvature, and the peak GCaMP signals, were higher for the cells in the preferred direction, these findings may not provide a complete explanation for the direction selective responses. For instance, evidence for possible differences in adaptation mechanisms is found in our sustained recordings on the tracking microscope which revealed a higher baseline calcium level in neurons that were responding to prolonged bouts of movement in the preferred direction.

A second possibility would invoke a circuit mechanism that involves inhibition. Our results have shown that the dendrite deformations observed in ddaE and ddaD occur at distinct phases of the segmental contraction cycle. During forward locomotion ddaE dendrites deform earlier than those of ddaD and the dendrites of ddaD deform earlier during reverse locomotion. Thus, the more strongly activated neuron is the first to experience deformation and it is possible that inhibition of the less strongly activated cell occurs during the delay. This model has similarities to the mechanisms that allow starburst amacrine cells to shape responses of direction selective ganglion cells of the vertebrate retina.

A third possibility is that dendrite deformations that progress in a distal to proximal direction are more strongly activating than those that progress in proximo-distal direction. Ionic currents that progress from distal to proximal might summate at a spike initiation zone reflected by calcium signals at the cell soma. In contrast, proximal to distal dendrite deformations would show reduced summation since the currents would progress in a direction that is moving away from the cell body. This model predicts passive dendrites in Class I neurons that lack strongly voltage gated currents.

Fourth, as with other mechanosensory systems the cellular transduction machinery of the Class I neurons may be constructed with an inherent asymmetry that causes it to be more sensitive to the forces that are generated in the preferred direction of movement. This model is appealing due to the involvement of the *Tmc* family of ion channels in the mechanically driven responses of both the Class I neurons and hair cells of the inner ear. Thus, elucidating the cellular ultrastructure of the *Tmc* dependent transduction machinery of Class I neurons will be a fascinating subject for future study.

Finally, our results indicate that the responses of the Class I neurons are consistent with the previously proposed “mission accomplished” model, but we add into this model the feature of direction selectivity. The highest responses of the neurons coincide with the phase of the segmental contraction cycle when the muscles of the segment are most fully contracted (ie. mission accomplished). The timing of this peak Class I response may facilitate the progression of the wave of neural activity in the larval ganglion to initiate contraction of the next segment, and the signals may also help to terminate the contraction of the preceding segment and within the contracting segment of the traveling wave. It is noteworthy that neurons of the larval ganglion have been identified that show specific activity during bouts of forward locomotion and backwards locomotion respectively[22, 23]. As the larval connectome mapping proceeds, it will be interesting to determine how sensory input from ddaE and ddaD impacts CNS circuits that are specifically engaged during forward and backward locomotion respectively.

## Materials and Methods

### Fly Strains and Husbandry

The following fly strains were used: *w;UAS-GCaMP6f;2-21-GAL4, w;UAS-GCaMP6f; 2-21-GAL4 Tmc*^1^, *w; UAS-6xmCherry, yw;UAS-mCD8::GFP; 2-21-GAL4*

Flies used in Figures 1,2,4 and 5 were maintained on Bloomington cornmeal medium. Larvae in these figures were collected from yeasted apple juice plates 24-48 hours after egg laying.

### Confocal Microscopy

L1 larvae were placed into a water filled channel (150 x 150 μm) through 3% agarose. Larvae were manually size selected under a dissecting microscope so that they fit snugly into the channels. Friction from the sides of the agarose channels slows the larvae as they crawl thus facilitating the analyses. Fluorescent signals emitted from GCaMP6f were recorded through a number 1.5 coverslip (forming the fourth side of the channel) on the stage of an inverted Zeiss LSM V Live confocal microscope that was equipped with a Physik Instrumente P-725 Piezo focus drive and a 40x 1.3NA PLAN-NEOFLUAR lens. Excitation was with a 488nm laser and emission was collected through a LP 505 filter. Image series were taken at zoom 1.5 with XY pixel dimensions ranging from 512×196 to 512×170. Volumes consisted of 10-12 Z slices and were collected at 9-15 volumes per second.

### Image Analyses

For confocal GCaMP6f experiments, maximum intensity projections (MIP) of the Z-volume time series were generated by the Zeiss Zen software package. Neuronal cell bodies were then tracked through the MIP time series using custom scripts in Matlab with user curation to ensure accurate tracking (see Supplemental Video 7 for an exemplar of ddaE tracking). The percentage change of GCaMP6f fluorescence intensity over time was calculated as ΔF/F for each image of the time series as [F(t) - F(B)]/F(B), where F(t) is the GCaMP6f fluorescence intensity of a ROI drawn around the cell soma of interest in each image, and F(B) is the basal GCaMP6f fluorescence intensity, which was determined by the basal GCaMP6f fluorescence intensity when the larvae were stationary.

The segmental contraction cycle was defined by the distance (D) between the pairs of neurons (ddaE or ddaD) in neighboring segments. The distance between cells during locomotion changes with a cycle of repetitive shortening and lengthening. A cycle starts when the distance between the neuron-pair is at a maximal distance which then reaches a minimum and then returns back to maximum again. We plotted the distance between the centroids of neighboring neuron-pairs over time were transformed to phase angles on a 360 degree cycle as follows: Theta_t_=(D_max_-D_t_)×(pi/(D_max_-D_min_)) gives the phase angle for the shortening phase of the cycle and the formula Theta_t_ =2*pi-(D_max_-D_t_)*(pi/(D_max_-D_min_)) gives the phase for the relaxation period where D_max_ is the maximal distance between the neuron-pair and D_min_ is the minimal distance between neuron-pair, and D_t_ is the distance between neuron-pair at a given time in the cycle.

Even with the rapid scanning procedure that we employed, the three-dimensional (3D) reconstruction of the dendritic arbors from moving animals posed challenges. For instance, during the time points with the most rapid movement, the dendritic field of the neuron was moving in the X, Y and Z axes during acquisition of Z-stacks. This caused each Z-section to be an individual snapshot of a portion of the moving arbor, and this movement resulted in mis-registration of slices in Z (visualized as a blurred image of the arbor when the stacks were displayed as a maximum intensity projection). Thus, the models of dendritic architecture shown in Figure 1 and 2 relied upon computational approaches. Mis-registration was minimized using an algorithm that realigned the slices in the Z axis based on the image intensity using the free-form deformation (FFD) technique[24]. However, the low axial resolution (sampling) in the Z-axis, along with the high degrees of freedom of the dendritic morphology, render most automated tracking methods, including ours in [25], unreliable. Therefore, the realigned dendrites were analyzed using a semi-automatic Computer Vision framework for 3D neurite tracing (in each image stack) and tracking (over time), considering a piece-wise articulated neuron “accordion” model.

Specifically, the extracted dendritic traces from non-moving, stationary time points allow us to generate a “ground truth” (prototype) model of the dendritic morphology (“average” morphology) for a particular neuron of interest. This ground truth morphology was projected on a surface that we call an “accordion” model: it consists of interconnected rectangular planes/strips, whose longer side is normal to the XY orientation of movement (along the agarose channel) (Fig. S3, Supplemental Video 12). Each strip is the linear spatial approximation of where the local maximum average intensity path of a dendrite branch is located (in a linear regression-like fashion). During motion, adjacent strips change orientation with respect to each other (and thus the “accordion” term), i.e., the pair-wise joint angles change, to approximate the local branch deformations (Fig. S3). This helps reduce the degrees of freedom and most importantly resolves ambiguities due to low resolution along the Z-axis. At each instance, the neuron accordion model is warped *in silico*, piece-wise for each strip, by finding the best-fit configuration to the recorded confocal data, which in turn deforms locally the corresponding ground truth dendrites to generate 3D estimates of the dendritic morphology. Our framework finds the best accordion (articulation) configuration for the neuron using the simulated annealing technique [26], based on user interaction, image features, and local shape. Users provide the X-coordinate (direction along the agarose channel) of the farthest tips of dendrites to help handle dendrite detection issues (Fig. S3). Local shape acts as regularization term to ensure smoothness of spatio-temporal transformations, penalizing non-planar configurations, i.e., enforcing angles between adjacent strips near 180 degrees. From the neuron accordion model, we were also able to estimate the dendrite curvature is using the sum of the angles between adjacent strips.

### 2P imaging of neuronal activity in freely behaving larvae

*w;UAS-GCaMP6f;2-21-GAL4* was crossed to *w; 20XUAS-GCaMP6f; 20XUAS-6XmCherry-HA* (created from Bloomington stock #42747 and #52268 using #3704 balancer). Mated flies were allowed to lay eggs for 24 hours at 25°C on cornmeal-based food. F1 progeny at second instar larval stage, 48-72 hr AEL, were separated from the food with 30% sucrose solution and washed in water. Larvae were hand selected for size.

The two-photon tracking microscope is described in detail in Karagyozov, Mihovilovic-Skanata, et al. 2018[9]. Briefly, ultrashort pulses with a central wavelength of 990 nm are focused through a 40X/1.15NA objective onto a targeted neuron, where they excite both GCaMP6f and hexameric mCherry. Emitted photons are spectrally separated (green for GCaMP6f and red for mCherry) and collected by two PMTs. The focal spot is rastered in a cylinder around and through the neuron, and the rate of photon emission is decoded by a FPGA to update the position of a neuron in real time. The ratio of green to red fluorescence reported the neurons’ calcium dynamics.

For a single neuron, the position and activity were updated continuously at a rate of 2.8 kHz. To simultaneously track the cell bodies of ddaE and ddaD neurons, our tracking algorithm first scanned the focal spot 4 complete cycles (1.4 ms) around the cell body of ddaE neuron before moving to track the ddaD neuron for 4 cycles. Transitioning between the cell bodies required 0.7 ms of downtime during which neither neuron was tracked. This means we recorded the fluorescence and the position of each neuron for 1.4 ms out of every 4.2ms.

At the beginning of each experiment, we located and identified the neuron(s) to track using epifluorescence microscopy. To accomplish this, we temporarily immobilized the larva in the chamber using vacuum compression. After locking the tracker onto the targeted neuron(s), we slowly released the vacuum to allow free crawling. Following release, we did not further manipulate the pressure in the chamber.

Semi-automated tracking recorded the tail position of the larva in each video frame. We then measured the velocity of the neuron relative to the axis between the tail and the neuron. Positive velocity indicates forward movement, and negative velocity indicates reverse movement. We then segmented forward and reverse "bouts:" points where the velocity in one direction or another was maximal and the surrounding time intervals.

Since the larva was allowed to freely roam about a circular chamber, we were able to record activity of ddaD and ddaE neurons during larval body bends. Body bends were determined by visual inspection of raw behavioral videos, blind to the recorded neural activity, using imageJ. Activity during a body bend was measured as the baseline corrected ratio of green to red fluorescence during the frame with maximal body bend.

For each body bend, we recorded the bend angle, if the larva bent its body towards(ipsi) or away from the tracked neuron(contra), and the direction of its movement before and after the turn. Groupings: *all* – all sampled body bends of the given type; *forward* – all sampled body bends in which the larva was crawling forward before and after the bend; *backward* – all sampled body bends in which the larva was crawling backward before and after the bend; *simultaneous recording* ***–*** subset of bends measured when tracking both ddaD and ddaE simultaneously. Note that the total number of bends is greater than the sum of the number of “forward” and “backward” bends, because many bends accompanied a change in direction.

P-values were computed using student’s two sample t-test and represent the probability of observing a difference in mean ipsi and contra activity at least as extreme under the null hypothesis that both ipsi and contra activity are drawn from the same normal distribution.

---

**Supplementary Table 1.** Quantification of Larval Bends During Turning Behavior (Related to Figure 4)

|  |  |  |
| --- | --- | --- |
| **all** | n bends | p-value comparing ipsi to contra |
| ddaE ipsi | 32 | 0.61 |
| ddaE contra | 33 |  |
| ddaD ipsi | 29 | 8.1*10<sup>-8</sup> |
| ddaD contra | 33 |  |
| **forward** | n bends | p-value comparing ipsi to contra |
| ddaE ipsi | 14 | 0.70 |
| ddaE contra | 19 |  |
| ddaD ipsi | 9 | 2.2*10<sup>-6</sup> |
| ddaD contra | 18 |  |
| **backward** | n bends | p-value comparing ipsi to contra |
| ddaE ipsi | 8 | 0.27 |
| ddaE contra | 7 |  |
| ddaD ipsi | 12 | 0.06 |
| ddaD contra | 3 |  |
| **simultaneous recording** | n bends | p-value comparing ipsi to contra |
| ddaE ipsi | 11 | 0.85 |
| ddaE contra | 15 |  |
| ddaD ipsi | 11 | 8.5*10<sup>-6</sup> |
| ddaD contra | 15 |  |

**FigureS1 related to Figure 3:**
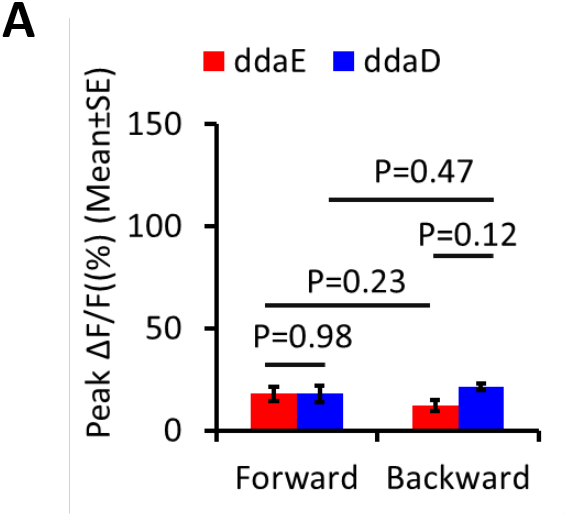
mCD8::GFP shows relatively minor movement related calcium response of *ddaE and ddaD* in both forwards and backwards locomotion A) Comparison of peak ΔF/F in ddaE and ddaD during larval forward and backward locomotion. There is no significant differences in peak ΔF/F between ddaE and ddaD during larval either forward(P=0.98) or backward locomotion(P=0.12). There is no significant differences in peak ΔF/F related to movement between larval forward and backward locomotion for ddaE(P=0.23), the same result for ddaD(P=0.47). Student’s t-test. n=12 (forward), n=3 (backward), n represents the number of movements examined.

**FigureS2 related to Figures 1,2,3,5,6,7:**
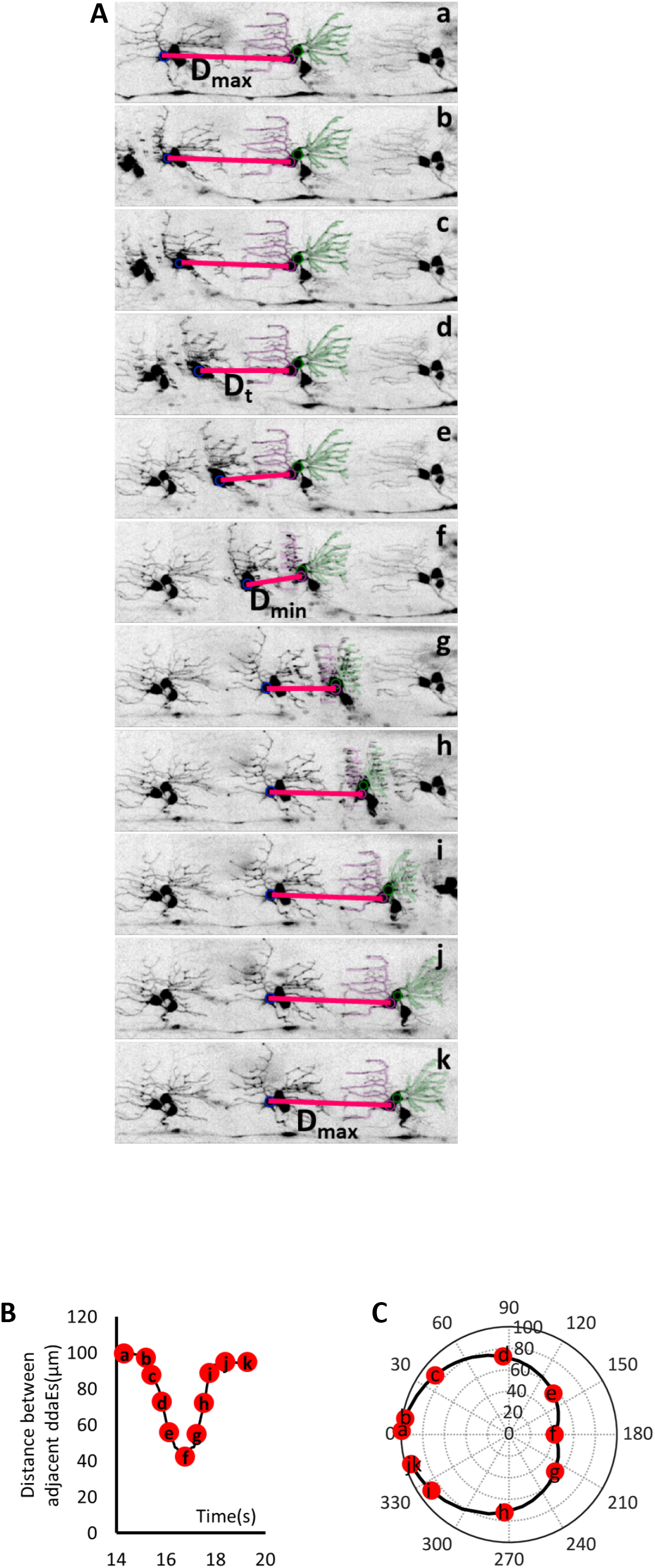
Definition of muscle contraction cycle. A) Real time images shows a muscle contraction cycle. The muscle contraction cycle is defined as the repetitive shortening and lengthening of the distance between the adjacent neuron-pair (ddaE or ddaD) in neighboring segments. A typical cycle starts from when the distance between the neuron-pair is at the maximal distance, reaches minimal, and back to maximal again as depicted in the A. B) Centroid distance between adjacent ddaE neurons was plotted as a function of time. Time for a contraction cycle is different for each movement or larva. C) Centroid distances of neighboring ddaE-pairs over time were plotted in phase of a 360 degree cycle. Position along the radial axis represents the distance between adjacent ddaE neurons showed in A. The centroid distances of neighboring neuron-pairs over time were further converted to the phase angles. For each cycle, maximal distance between neighboring neuron-pair (initiation of the contraction) is set to phase 0, and the minimal distance (initiation of relaxation) is set to phase 180. The end of the cycle is when it reaches the maximal distance (phase0/360). The formula Theta_t_=(D_max_-D_t_)*(pi/(D_max_-D_min_)) was used to convert distance to phase angle coordinate (0-180o), and Theta_t_ =2*pi-(D_max_-D_t_)*(pi/(D_max_-D_min_)) was used to convert distance to phase angle coordinate(181o to 360o), where Dmax is the maximal distances between neuron-pair, Dmin is the minimal distance between neuron-pair, and Dt is the distance between neuron-pair at a given time in the cycle.

**FigureS3 related to Figure 5.**
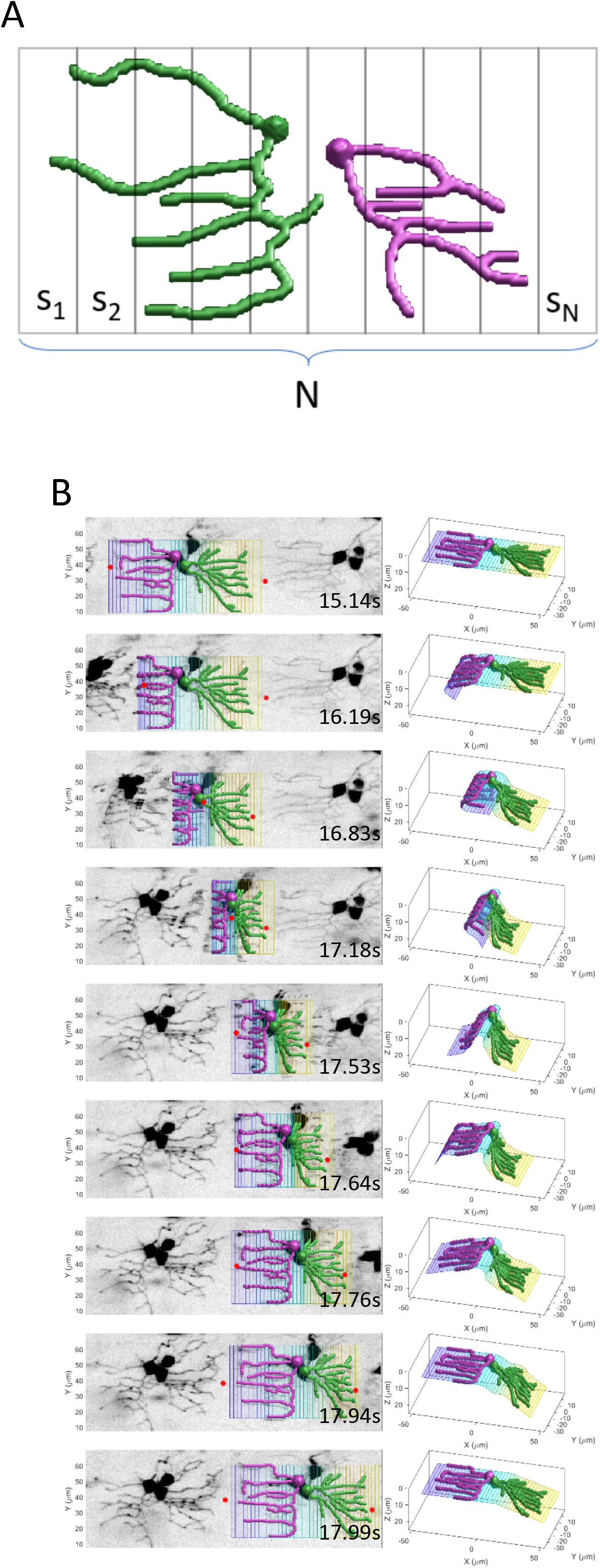
A) The neuron plane model enclosing the body segment consist of N strips S_1_, S_2_, S_3_, ….S_N_ B) The algorithm requires the X-coordinates of the farthest dendrite tips as shown by red points. ddaD and ddaE neurons are shown in green and magenta colors respectively. The colored strips constitute the neuron accordion model. (Left) the 2D view. (Right) the 3D view. From top to bottom, the neuron accordion model over a contraction cycle — the ground truth dendrites, and the deformed neurons in frames at various time points. Notes: ventral is up and anterior is to the right.

**FigureS4 related to Figure 4.**
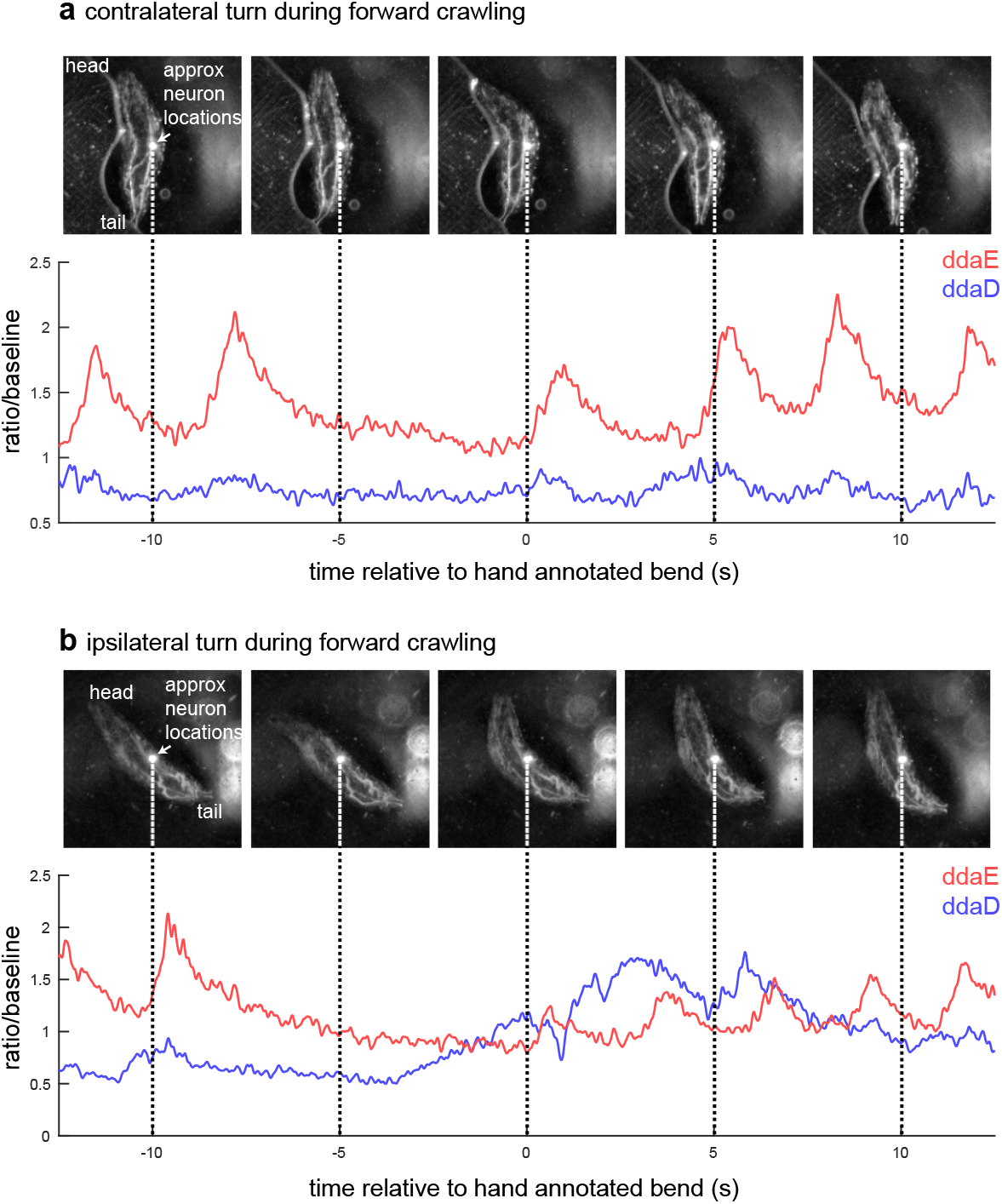
Representative images of body bends observed in the 2-photon tracking microscope. The top images in a and b are still images from video sequences of larval locomotion in the chamber. The bright spot in each panel indicates focal point of the 2-photon laser. The bottom shows the ratiometric traces of neural activity with dashed lines that indicate the time point at which the images have been extracted. **a)** Representative contralateral bend in which the larva turns towards the side of the body opposite to the tracked neurons. Note that ddaD remains quiescent. **b)** Representative ipsilateral bend which shows an increase in ddaD activity during the bend.

## Legends for Supplemental Videos

**Video S1 (Related to Figure 1):** Forward locomotion recording of larva expressing mCD8::GFP in the under control of the driver 221-GAL4. Scale bar is 40μm.

**Video S2 (Related to Figure 1):** Video of the 3D model for ddaE (magenta) and ddaD dendrites in forward locomotion.

**Video S3 (Related to Figure 2):** Backward locomotion recording of larva expressing mCD8::GFP in the under control of the driver 221-GAL4. Scale bar is 40μm.

**Video S4 (Related to Figure 2):** Video of the 3D model for ddaE (magenta) and ddaD dendrites in backward locomotion.

**Video S5 (Related to Figure 1):** Recording of GCaMP6F during forward locomotion. The GCaMP6F fluorescence in dendrites and soma of ddaE increases in response to larval forward movement, while ddaD fluorescence remains relatively stable during larval forward movement. Color map from blue to red denotes the activity from low to high.

**Video S6 (Related to Figure 2):** Recording of GCaMP6F during forward locomotion. The GCaMP6F fluorescence in dendrites and soma of ddaD increases during backward movement, while that of dendrites and some of ddaE remains relatively stable during larval backward movement. Color map from blue to red denotes the activity from low to high.

**Video S7 (Related to Figure 3):** GCaMP6F fluorescence recording of 3 bouts of backward and 3 bouts of forward locomotion. This recording also demonstrates the neuronal tracking software for the ddaE neuron. This larva first crawled backward 3 times and then moved forward 3 times. Color map from blue to red denotes the activity from low to high. Scale bar is 40μm. Note that video resolution is substantially reduced relative to the raw data to aid with download. Original data available upon request.

**Video S8 (Related to Figures 1, 2 and 3):** Recording of mCD8::GFP fluorescence in ddaE and ddaD neurons during backward locomotion. Only minor movement-related GFP changes in *ddaE* and *ddaD* are observed in larvae expressing mCD8::GFP in Class I da neurons.

**Video S9 (Related to Figure 4):** Infrared recording of larval behavior during period of time shown in Figure 4. The bright spot on the larva’s body indicates the excitation laser and indicates the approximate location of the neurons being tracked. Because feedback tracking keeps the neuron centered under the objective (a portion of which can be seen out of focus in the background of the video), it can be difficult to visualize the larva’s forward or backward progress. We have overlaid small cyan dots to show the fixed reference frame of the microfluidic device in which the larva is crawling. The upper left overlay shows activity of ddaD and ddaE for the previous 20 seconds, with the dashed line indicating the time of the current frame. The time in the lower left corresponds to the x-axis in Figure 4. Video was recorded at 10 fps. At the default playback rate of 30 fps, video is played 3x real speed.

**Video S10 (Related to Figure 6 and 7):** Recording of GCaMP6F during forward locomotion in *Tmc^1^* mutant larvae. No increase of GCaMP6F fluorescence in dendrites and soma of ddaE was observed in *Tmc^1^* mutants during forward locomotion. Color map from blue to red denotes the activity from low to high.

**Video S11 (related to Figures 6 and 7):** Recording of GCaMP6F during backward locomotion in *Tmc^1^* mutant larvae. No increase of GCaMP6F fluorescence in dendrites and soma of ddaD was observed in *Tmc^1^* mutants during backward locomotion. Color map from blue to red denotes the activity from low to high.

**Video S12 (related to Figure 5):** Illustration of a run of our accordion model that was used to estimate dendrite curvature.

